## Supplementary for "The genomic landscape of metastatic castration-resistant prostate cancers reveals multiple distinct genotypes with potential clinical impact"

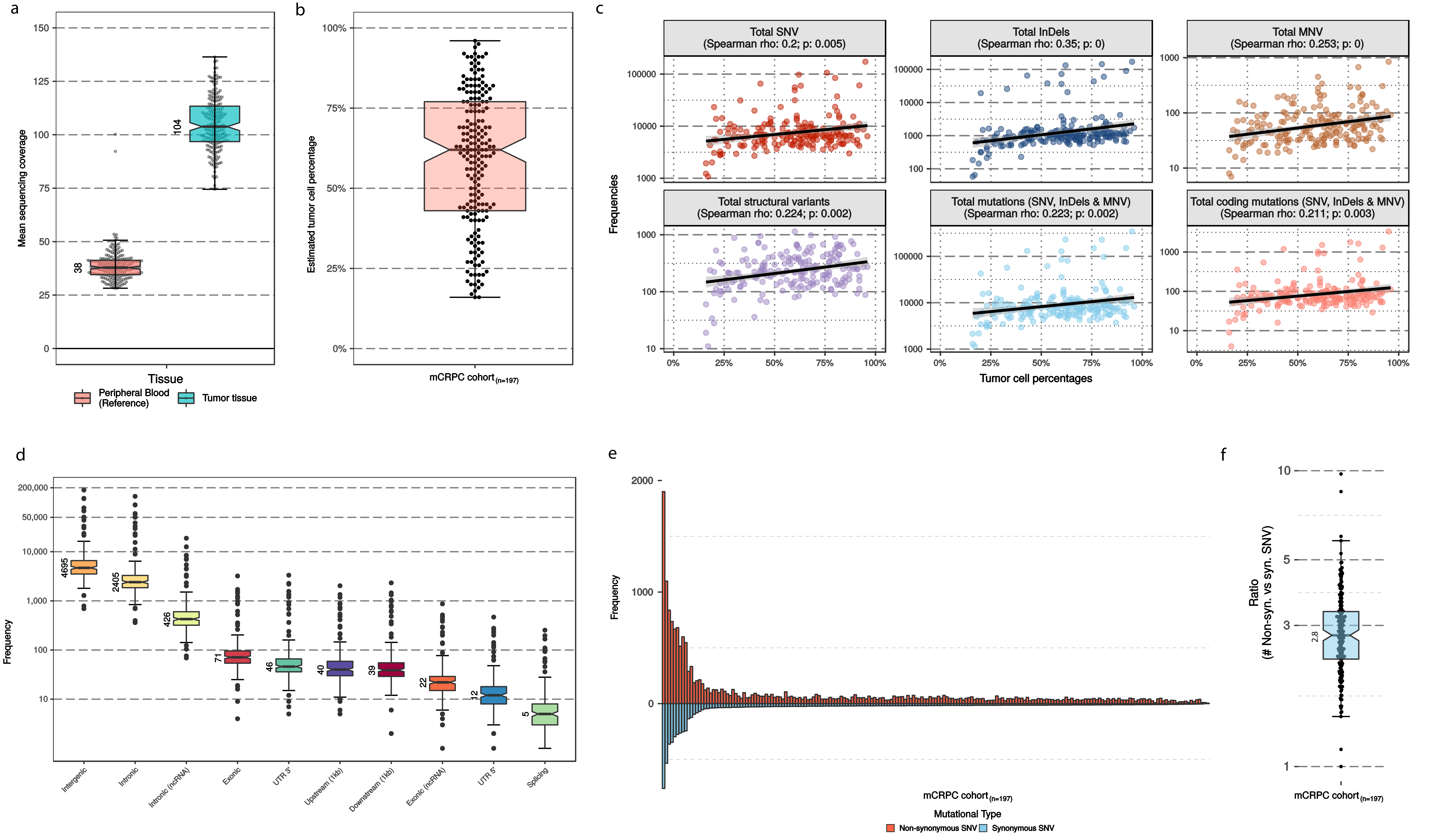

Supplementary figure 1 - Sequencing quality metrics
Overview of sequencing quality metrics.
(a) Bee-swarm boxplot with notch of the mean read coverage per sample of reference and tumor tissues. Boxplot depicts the upper and lower quartiles, with the median shown as a solid line; whiskers indicate 1.5 times the interquartile range (IQR). Data points outside the IQR are shown.
(b) Bee-swarm boxplot with notch of the estimated (in silico) cohort-wide tumor cell percentages. Boxplot depicts the upper and lower quartiles, with the median shown as a solid line; whiskers indicate 1.5 times the interquartile range (IQR). Data points outside the IQR are shown.
(c) Correlation (spearman) of estimated tumor cell percentages with observed aberrations per mutational category. Based on the low rank correlation coefficients (Spearman rho) we did not find high correlation with TC% and detected events, however a minor correlation could indeed be seen.
(d) Overview of the locations of variants (SNV / InDels / MNV) in respect to UCSC gene-models. Boxplot with notch depicts the upper and lower quartiles, with the median shown as a solid line; whiskers indicate 1.5 times the interquartile range (IQR). Data points outside the IQR are shown.
(e) Frequency of non-synonymous (red) and synonymous (blue) SNV per mCRPC sample.
(f) Ratio of non-synonymous over synonymous SNV for the entire mCRPC cohort. Bee-swarm boxplot with notch of the ratio. Boxplot depicts the upper and lower quartiles, with the median shown as a solid line; whiskers indicate 1.5 times the interquartile range (IQR). Data points outside the IQR are shown.

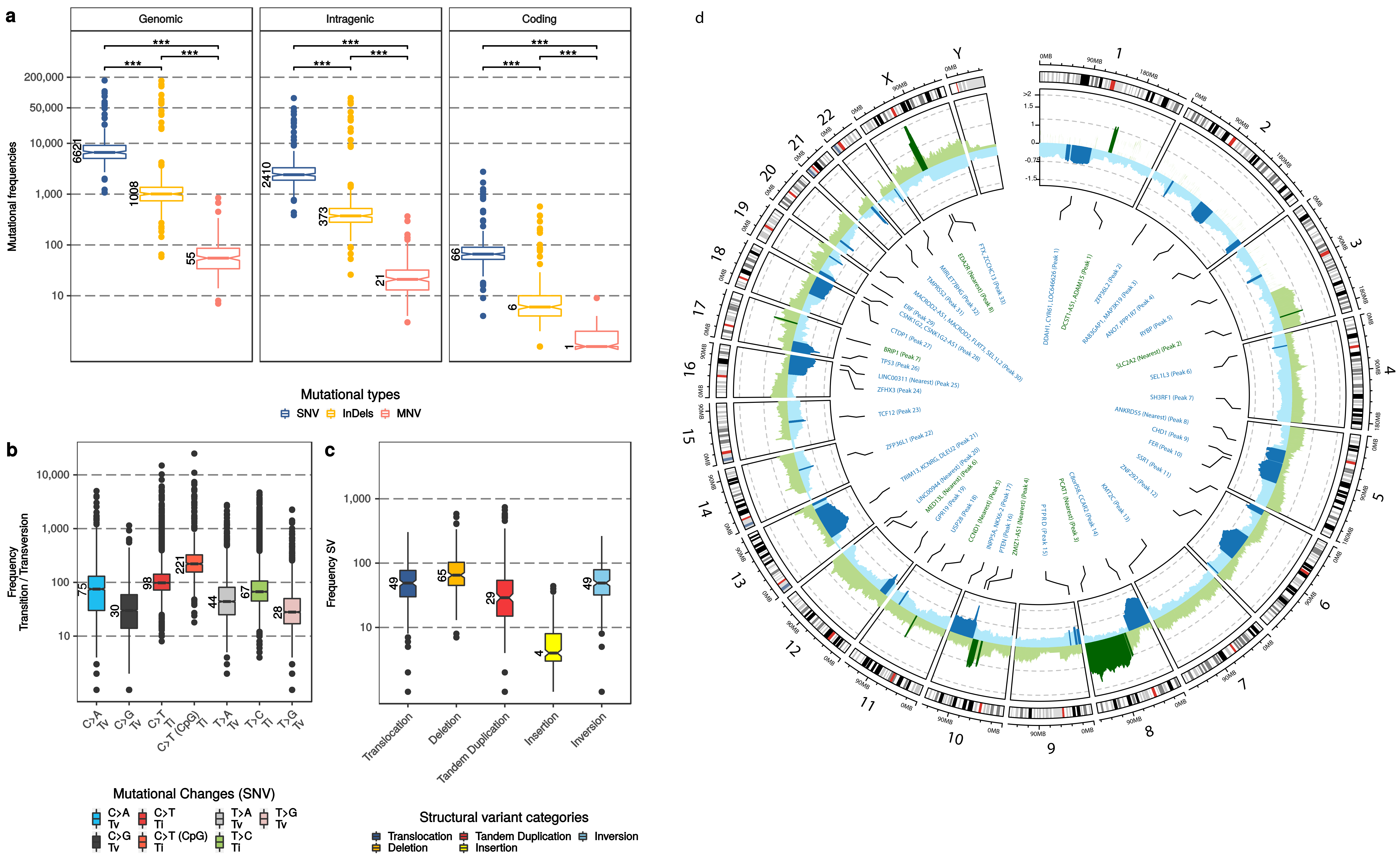

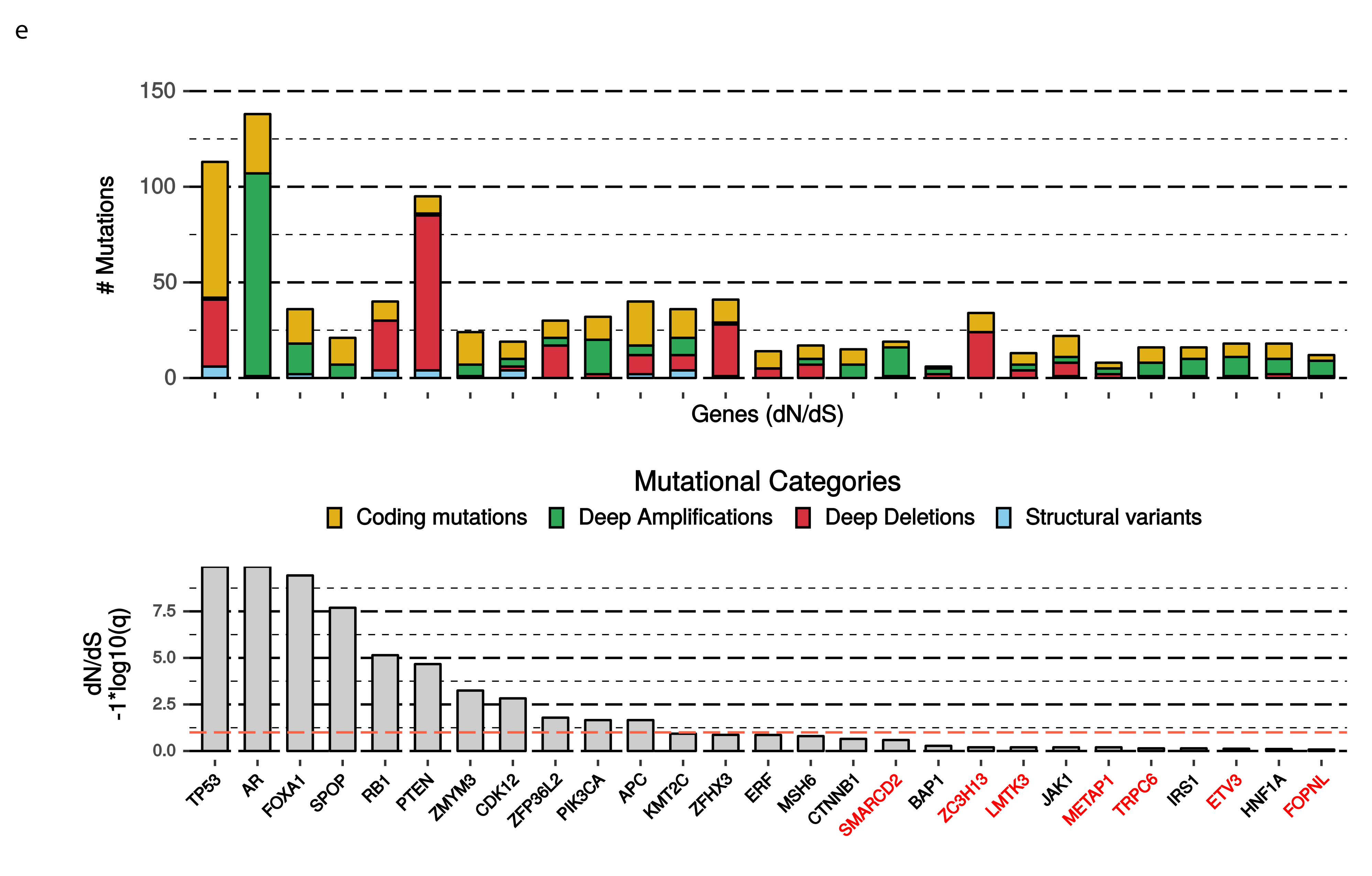

**Supplementary figure 2 - Overview of cohort-wide mCRPC somatic characteristics**

(a) Number of SNV (blue), InDels (yellow) and MNV (orange) per whole-genome sequenced sample over three resolutions; genome-wide, within intragenic regions and within coding regions. Boxplot with notch depicts the upper and lower quartiles, with the median shown as a solid line; whiskers indicate 1.5 times the interquartile range (IQR). Data points outside the IQR are shown. Statistical significance (Wilcoxon rank-sum test) is denoted per comparison.

(b) Type of genome-wide SNVs. Transition (Ti) and transversion (Tv), with a special attention for C to T Ti in CpG context, are indicated per sample. Boxplot with notch depicts the upper and lower quartiles, with the median shown as a solid line; whiskers indicate 1.5 times the interquartile range (IQR). Data points outside the IQR are shown.

(c) Frequency of Tandem Duplications (DUP), Insertions (INS), Inversions (INV), Deletions (DEL) and interchromosomal translocations (BND) are indicated per sample. Boxplot with notch depicts the upper and lower quartiles, with the median shown as a solid line; whiskers indicate 1.5 times the interquartile range (IQR). Data points outside the IQR are shown.

(d) Overview of recurrent copy number aberrations as detected by GISTIC2. G-scores are depicted on the y-axis ranging from 0 to ≥ 2. Regions with amplifications (G-score > 0) are depicted in green and deletions (G-score < 0) in blue. Regions with significant (and recurring) copy number aberrations (q ≤ 0.1) are denoted with a darker shade of green or blue, respective of amplification or deletion. Per region, the foci of maximal amplification or deletion (focal peaks; q ≤ 0.1) are denoted in the inner track; the peak identifier is also denoted as presented in supplementary table 3.

(e) Overview of genes detected by the dN/dS algorithm and corresponding mutational categories. Genes not present in one of our lists of known (onco)genes are colored red; (COSMIC v85, CGI, CIVIC and the list from Martincorena et al.). The upper figure displays absolute frequencies per mutational category in the detected genes and the lower figure displays the respective q-value (-1*log10(q)). The red line in the bottom figure indicates the threshold for statistical significance (q = 0.01).

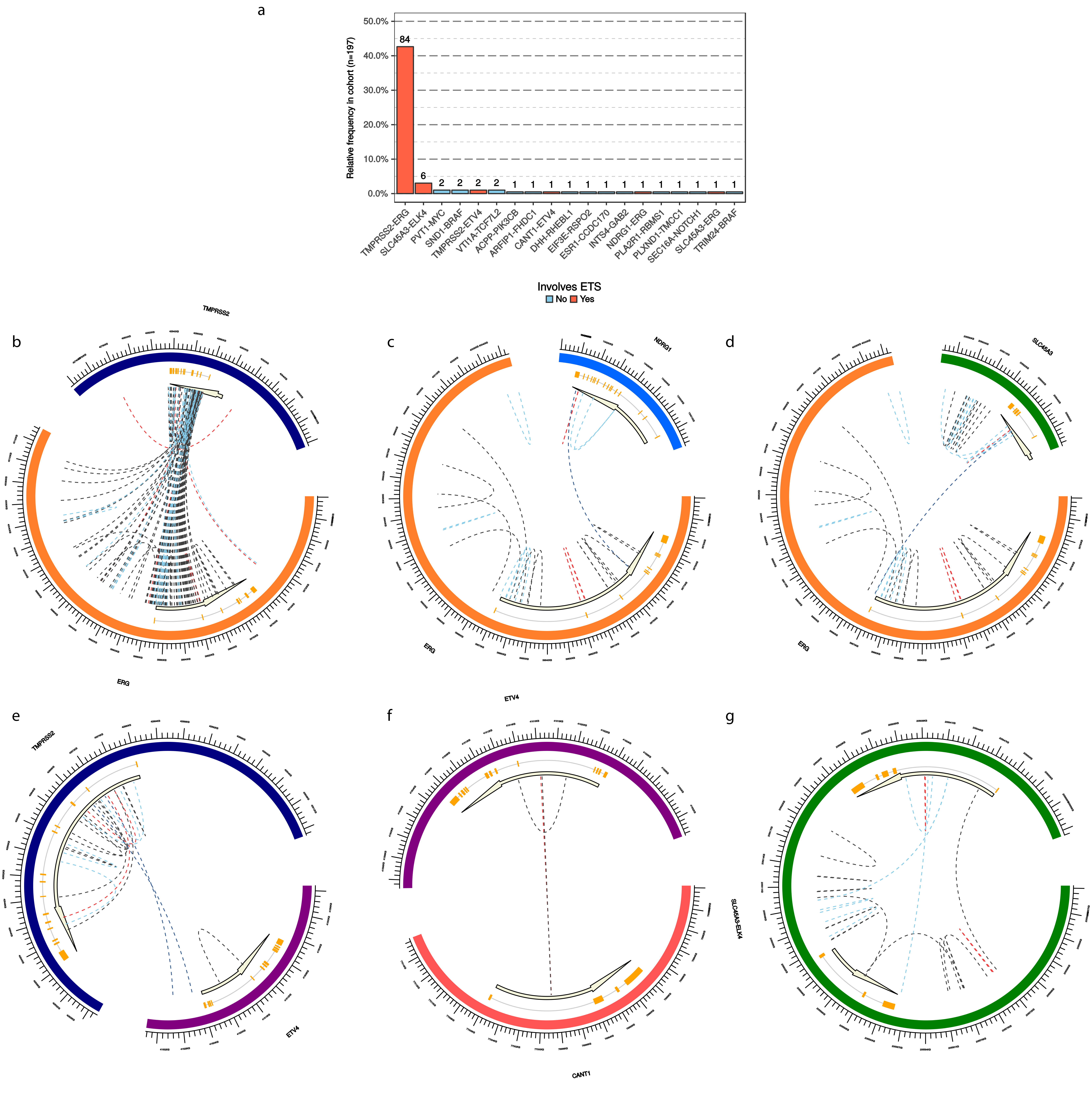

**Supplementary figure 3 - Overview of genomic (ETS) fusions**

(a) Relative frequency of observed genomic aberrations resulting in potential fusion products. Fusions involving ETS genes are depicted in red, other potential fusion partners in blue. Numbers above the bars indicate the absolute number of patients with the genomic aberration in the mCRPC cohort.

(b – h) Overview of structural variants involving the TMPRSS2 and ERG loci (b), NDRG1 and ERG loci (c), SLC45A3 and ERG loci (d), TMPRSS2 and ETV4 loci (e), CANT1 and ETV4 loci (f) and the SLC45A3 and ELK4 loci (g) in the mCRPC cohort. Interchromosomal translocations are colored in dark blue, deletions in black, insertions in yellow, inversion in light blue and tandem duplications in red. Orange boxes indicate exons; black line connecting the boxes are introns.

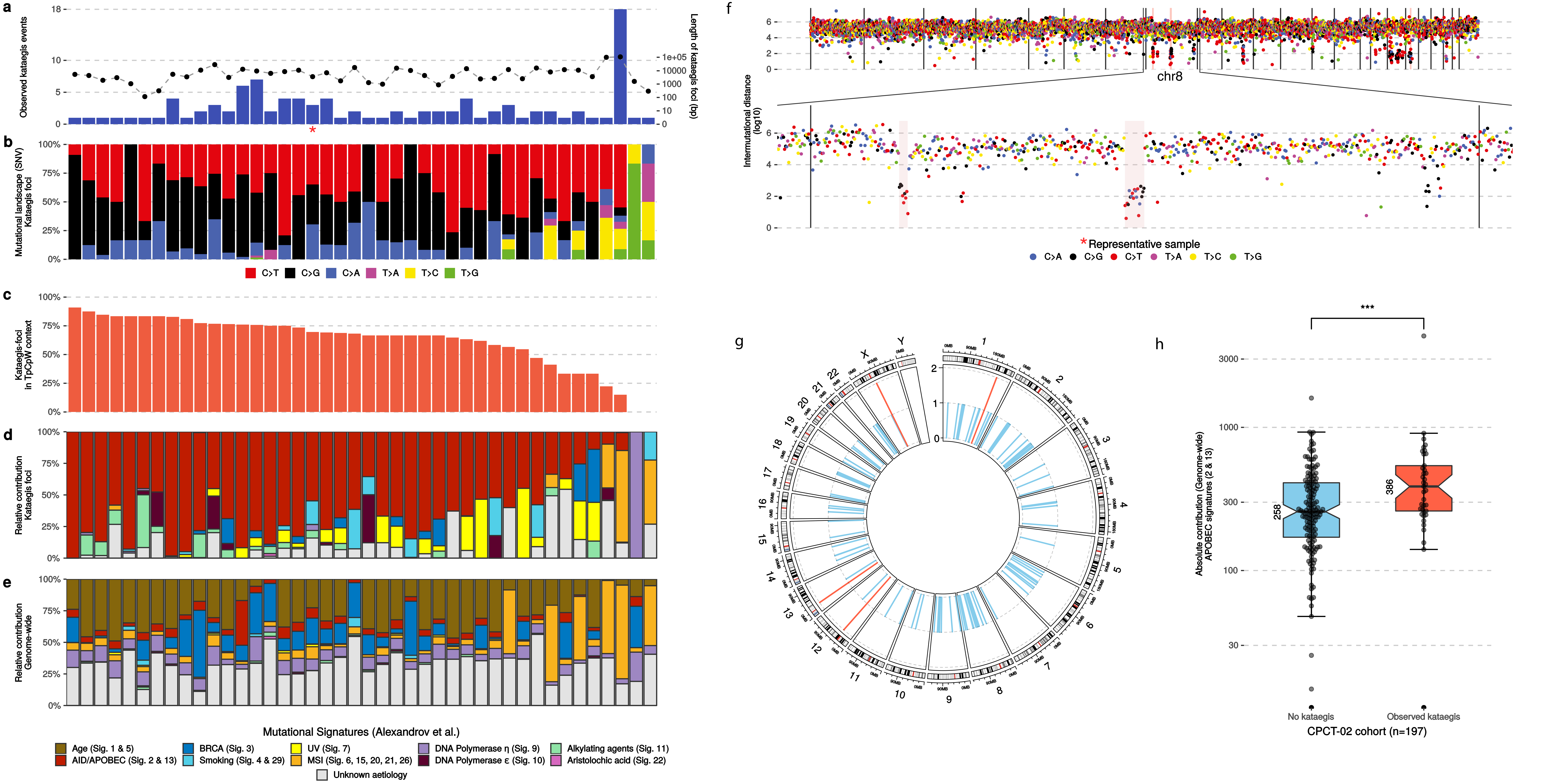

**Supplementary figure 4 - Kataegis prevalence in mCRPC**

(a) Number of observed kataegis events in mCRPC cohort samples (n = 42, blue bars) and the respective genomic width of all observed kataegis foci per sample (right y-axis; black points).

(b) Relative frequency of mutational contexts (of SNV) found in all observed kataegis foci per sample.

(c) Relative frequency of SNV in observed kataegis foci in APOBEC-related TpCpW mutational context. W stands for T or A.

(d) Relative contribution to mutational signatures (COSMIC) within the kataegis foci.

(e) Relative contribution to mutational signatures (COSMIC) of all genome-wide events of the sample.

(f) Representation of two distinct kataegis foci on chromosome 8 within a single respective sample (highlighted with * in a). SNV (colored on Ti/Tv type) are shown with relative genomic distances (in log10) to neighboring SNV. Observed kataegis foci are highlighted with a transparent red background.

(g) Frequency and locations of cohort-wide observed kataegis foci, binned per 1 Mbp. Bins with 2 kataegis events in distinct samples are colored red, else blue.

(g) Absolute contribution of APOBEC signatures (2 & 13) in samples without (n = 155) and with observed kataegis (n = 42). Bee-swarm boxplot with notch of the mean absolute contribution of APOBEC signatures (2 & 13). Boxplot depicts the upper and lower quartiles, with the median shown as a solid line; whiskers indicate 1.5 times the interquartile range (IQR). Data points outside the IQR are shown. Statistical significance was tested with Wilcoxon rank-sum test.

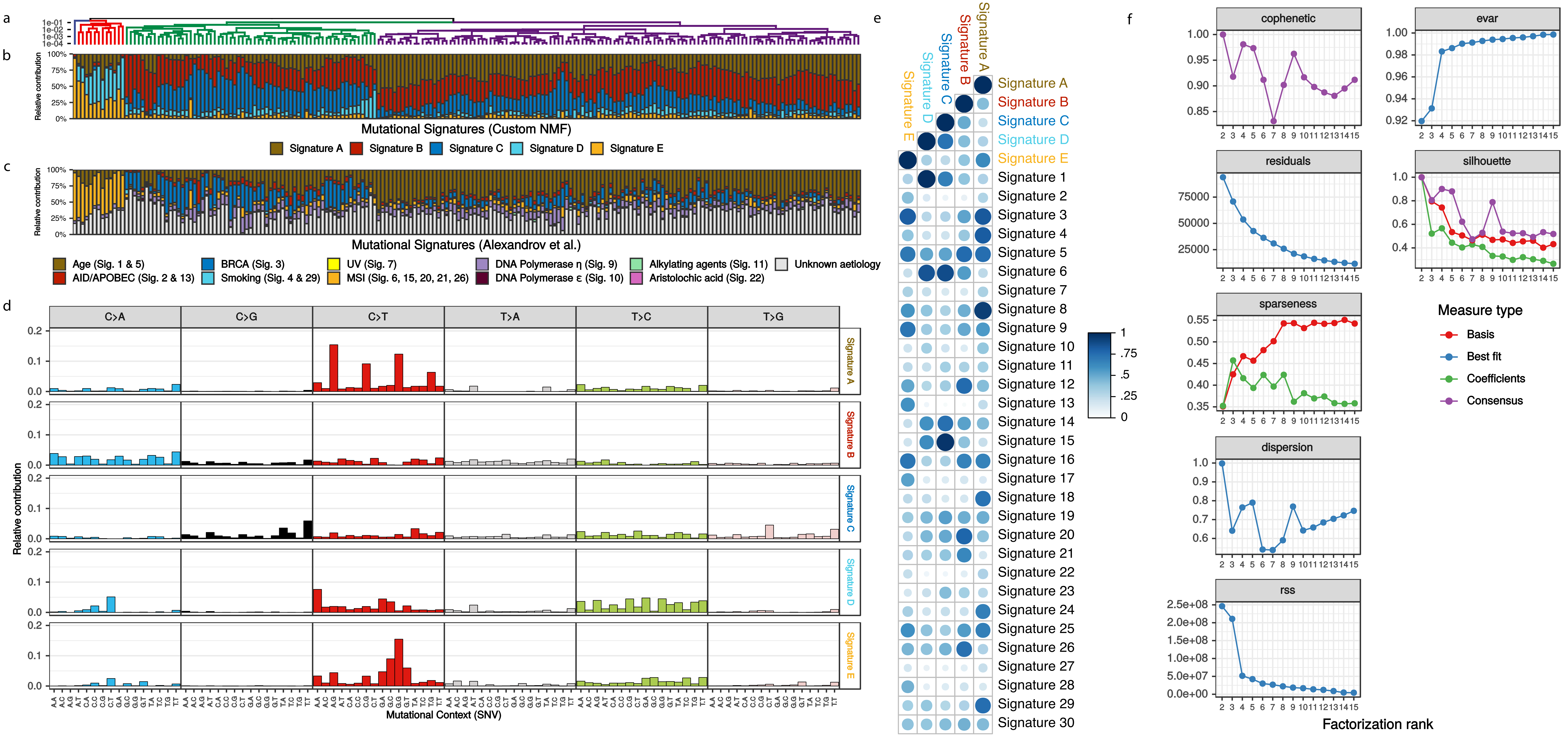

**Supplementary figure 5 - Mutational signature analysis**

(a) Dendrogram of unsupervised clustering (Euclidean distance; Ward.D method) on absolute contributions of SNVs in custom signatures A-E.

(b) Relative contribution to the five custom mutational signatures A-E.

(c) Relative contribution to COSMIC mutational signatures.

(d) Relative distribution of the 96 mutational contexts present in the custom signatures.

(e) Correlation (cosine similarity) of novel signatures with COSMIC signatures. The size of the dot reflects the cosine similarity, with higher cosine similarity values shown as larger dots. The color gradient indicates the level of cosine similarity.

(f) Quality metrics of NMF between two to fifteen ranks.

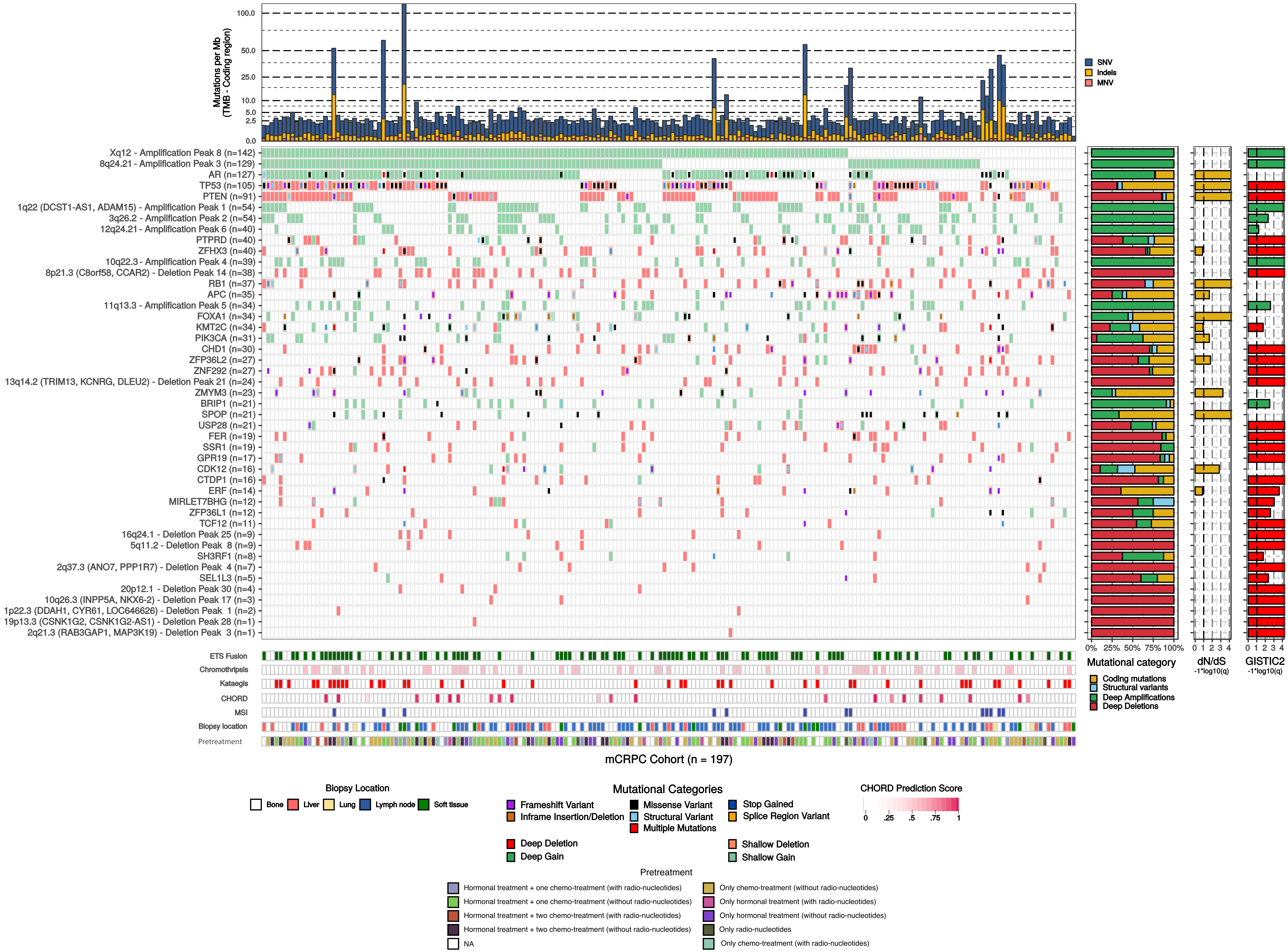

**Supplementary figure 6 - The mutational landscape of mCRPC seems unrelated to treatment history**

The upper track displays the number of genomic mutations per Mbp (TMB) of SNV (blue), InDels (yellow) and MNV (orange) categories. The heatmap displays the type of mutation(s) per sample; (light-)green or (light-)red backgrounds depict copy number aberrations whilst the inner square depicts the type of (coding) mutation(s). Relative proportions of mutational categories (coding mutations [SNV, InDels and MNV] (yellow), SV (blue), deep amplifications [high-level amplifications resulting in many additional copies] (green) and deep deletions [high-level losses resulting in (near) homozygous losses] (red)) per gene and foci are shown in the bar plot next to the heatmap. Narrow GISTIC2 peaks covering ≤ 3 genes were reduced to gene-level rows if one of these genes is present in the dN/dS (q ≤ 0.1) analysis or is a known oncogene or tumor-suppressor. For GISTIC2 peaks covering multiple genes, only deep amplifications and deep deletions are shown. Recurrent aberrant focal genomic foci in gene deserts are annotated with their nearest gene. Significance scores (-1*log10(q)) of the dN/dS and GISTIC2 analysis are shown on the outer-right bar plots; bars in the GISTIC2 significance plot are colored red if these foci were detected as a recurrent focal deletion and green if detected as a recurrent focal gain. Per sample, the presence of (predicted) ETS fusions (green), chromothripsis (light pink), kataegis (red), CHORD prediction score (HR-deficiency) (pink gradient), MSI status (dark blue), biopsy location and treatment history are shown as bottom tracks.

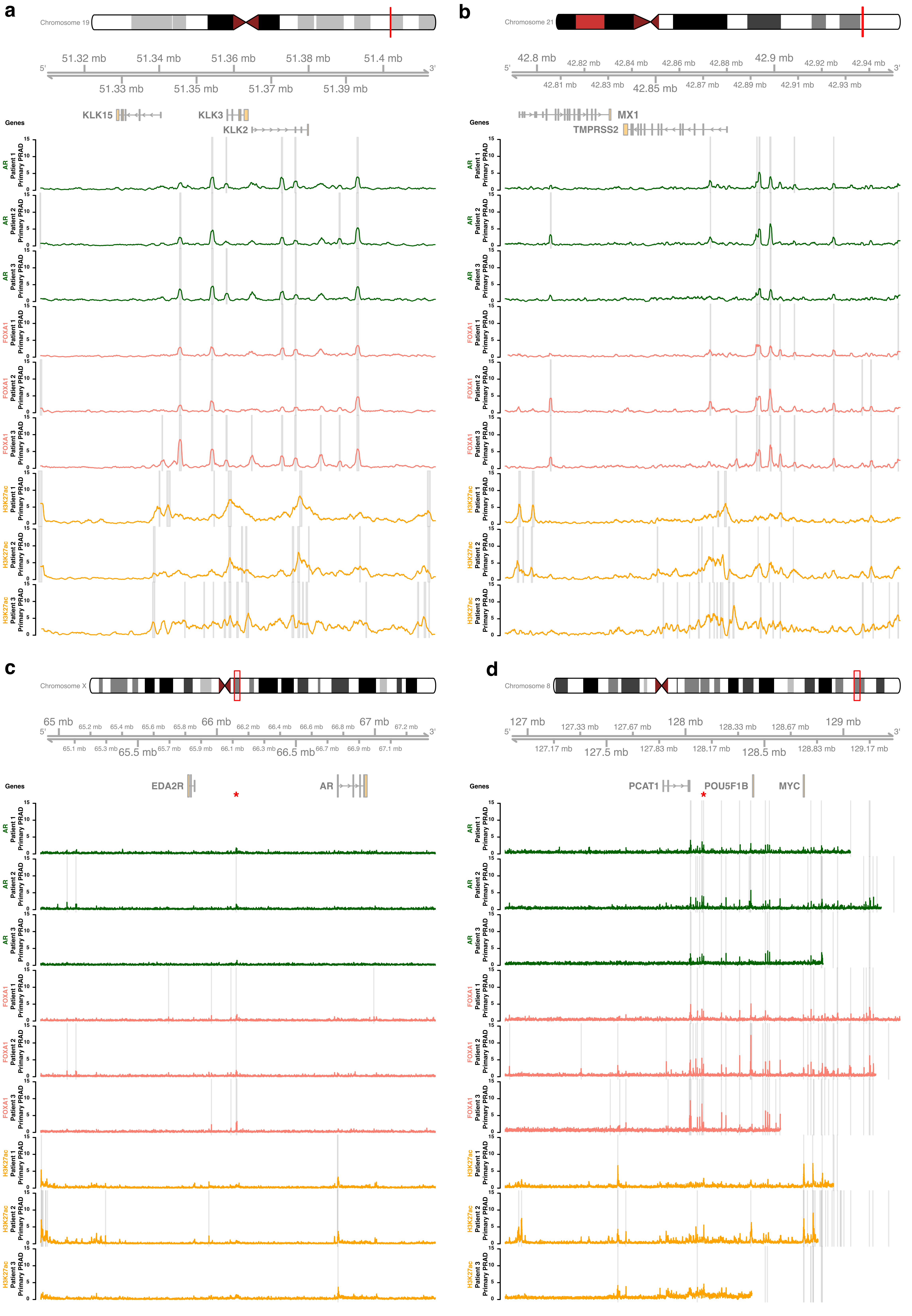

**Supplementary figure 7 - ChIP-seq profiles in primary prostate cancer for known driver genes**

ChIP-seq profiles from three independent primary prostate cancer patients surrounding the AR and PCAT1/MYC gene loci (with 1.25 additional Mbp up-/downstream) and two known AR-regulated positive controls (KLK3 and TMPRSS2 with additional 0.5 Mbp up-/downstream). Per subplot, the upper panel displays the selected genomic window and the overlapping genes. The 1th to 3th tracks represent AR ChIP-seq profiles (median read-coverage per 1000bp windows) in the three primary prostate cancer patients. The 4th to 6th tracks represent FOXA1 ChIP-seq profiles (median read-coverage per 1000bp windows) in the three primary prostate cancer patients. Finally, the 7th to 9th track represent H3K27ac ChIP-seq profiles (median read-coverage per 1000bp windows) in the three primary prostate cancer patients. ChIP-seq peaks (MACS/MACS2; q < 0.01) are shown as grey transparent lines per respective sample.

(a) ChIP-seq profiles surrounding the positive control KLK3 region.
(b) ChIP-seq profiles surrounding the positive control TMPRSS2 region.
(c) ChIP-seq profiles surrounding the AR region. The red asterisk denotes the location of the amplified region within the mCRPC setting.
(d) ChIP-seq profiles surrounding the PCAT1/MYC region. The red asterisk denotes the location of the amplified region within the mCRPC setting.

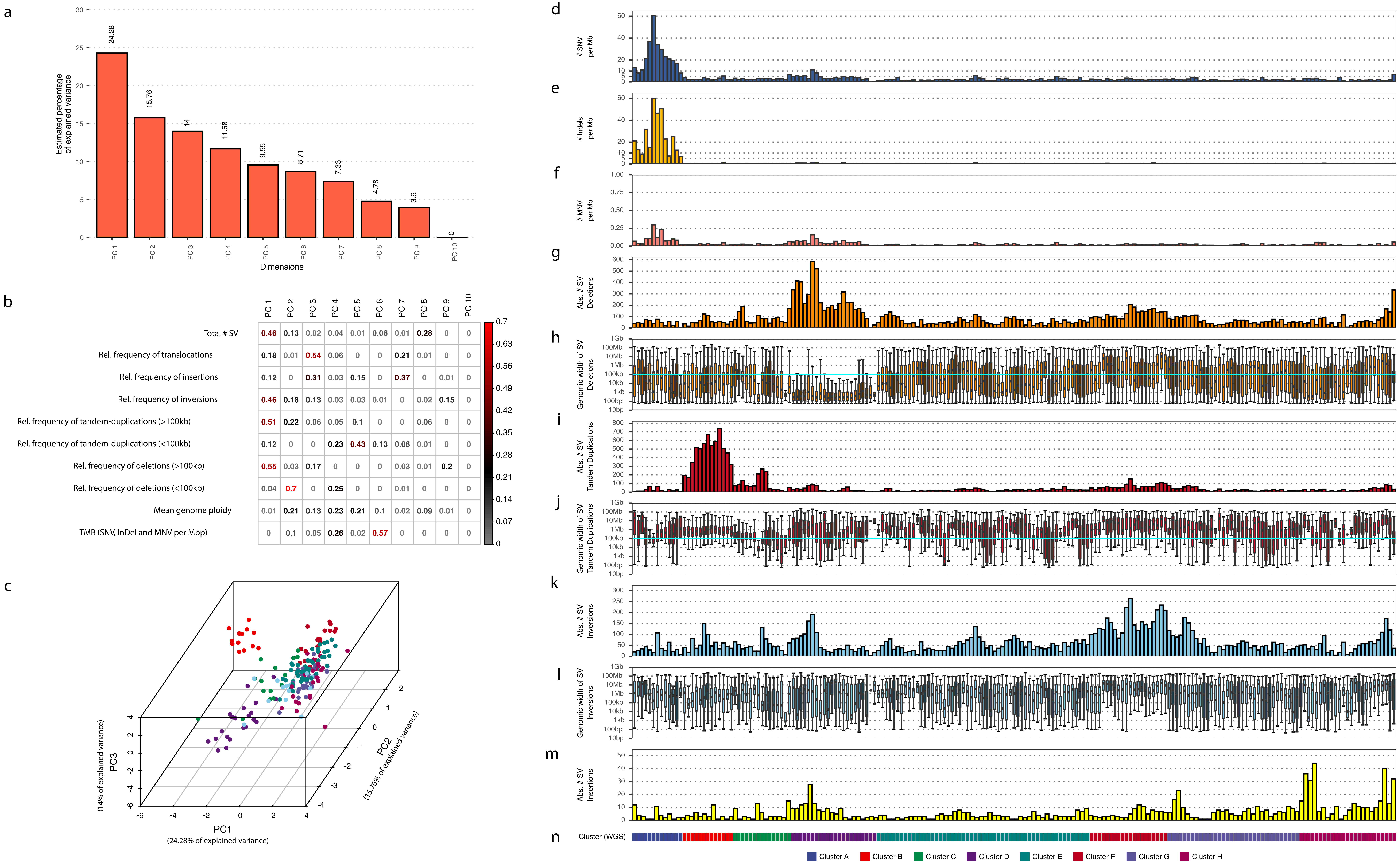

**Supplementary figure 8 - Rationale of the chosen genomic features for unsupervised clustering**

Principal component analysis (PCA) and overview of the genomic features included in the unsupervised clustering analysis highlighting the chosen size cut-offs and striking differences between samples.

(a) Overview of the explained variance per principal component (PC) in PCA.

(b) The quality of representation for each feature per principal component (cos2), this ranges from 0 (no importance / representation in PC) to 1 (absolute importance / representation in PC). Color gradient (0 to 0.7) denotes cos2, red values denote important / representation of feature within PC. Numbers shown are the cos2 values.

(c) Visualization of the first three principal components of PCA, each sample is colored based on their assigned cluster (depicted in n) after unsupervised clustering on their genomic features.

(d) All genome-wide somatic SNVs per Mbp (square root scale).

(e) All genome-wide somatic InDels per Mbp (square root scale).

(f) All genome-wide somatic MNV per Mbp (square root scale).

(g) Absolute number of deletions (SV) per sample.

(h) Distribution of the genomic width of deletions (SV) per sample. Cyan line indicates the chosen size cut-offs (< 100 kbp and ≥ 100 kb).

(i) Absolute number of tandem duplications (SV) per sample.

(j) Distribution of the genomic width of tandem duplications (SV) per sample. Cyan line indicates the chosen size cut-offs (< 100 kbp and ≥ 100 kb).

(k) Absolute number of inversions (SV) per sample.

(l) Distribution of the genomic width of inversions (SV) per sample.

(m) Absolute number of insertions (SV) per sample. Genomic width of insertions could not be estimated accurately due to repeat-like sequences.

(n) Assigned clusters based on unsupervised clustering of genomic features.

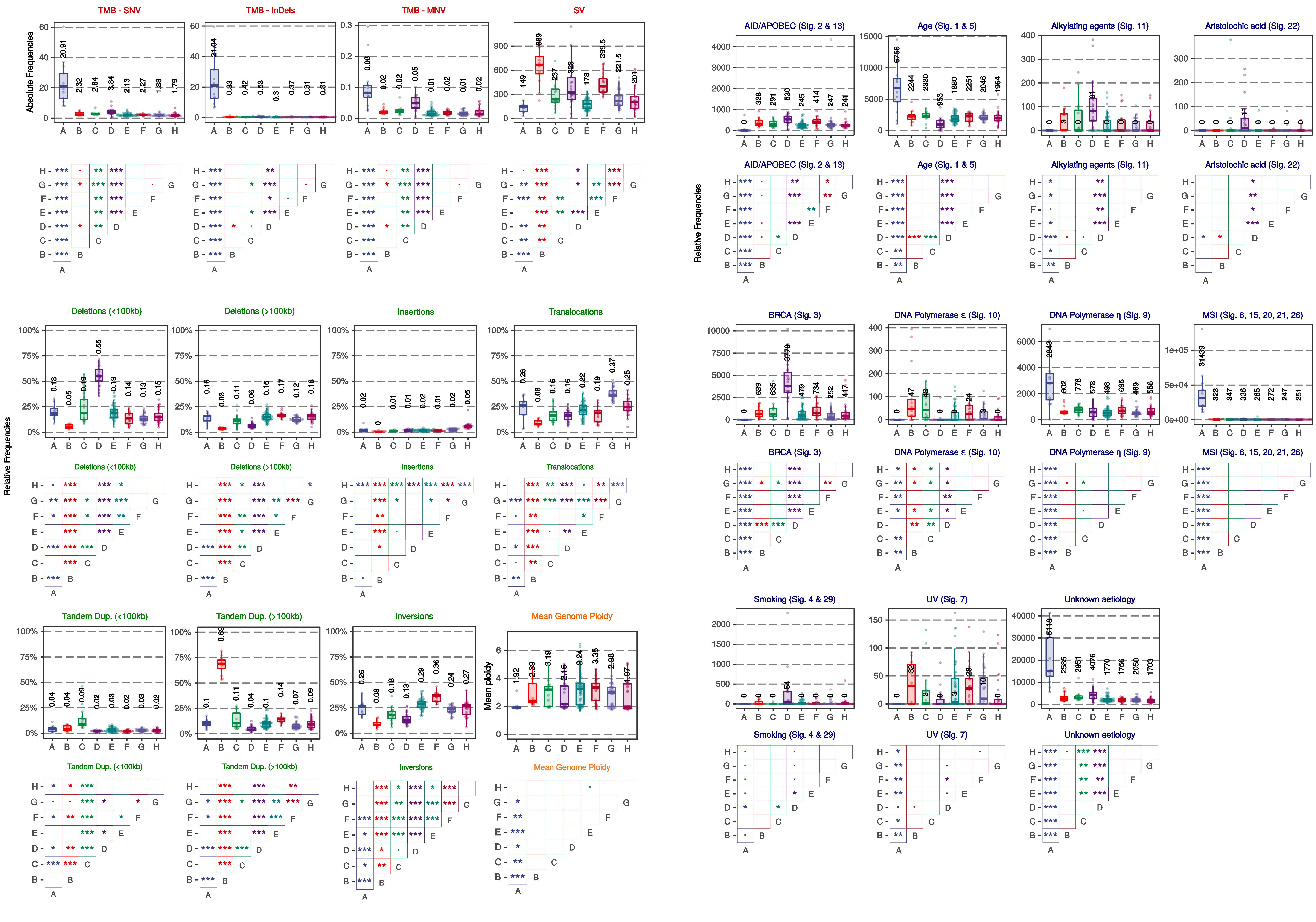

**Supplementary figure 9 - Cluster characteristics**

Overview of genomic characteristics and COSMIC mutational signatures per cluster (A-H) derived from unsupervised clustering of the mCRPC cohort using basic WGS characteristics. Bee-swarm boxplot depicts the upper and lower quartiles, with the median shown as a solid line; whiskers indicate 1.5 times the interquartile range (IQR). Data points outside the IQR are shown. A pairwise Wilcoxon rank-sum test (BH correction) was performed to detect statistically significant differences between clusters; * denotes p ≤ 0.05, ** denotes p ≤ 0.01, *** denotes p ≤ 0.001. Significant differences of events were also found in clusters without a clear biological association (C, E and G-H), such as increased numbers of translocations in cluster G and insertions in cluster H.

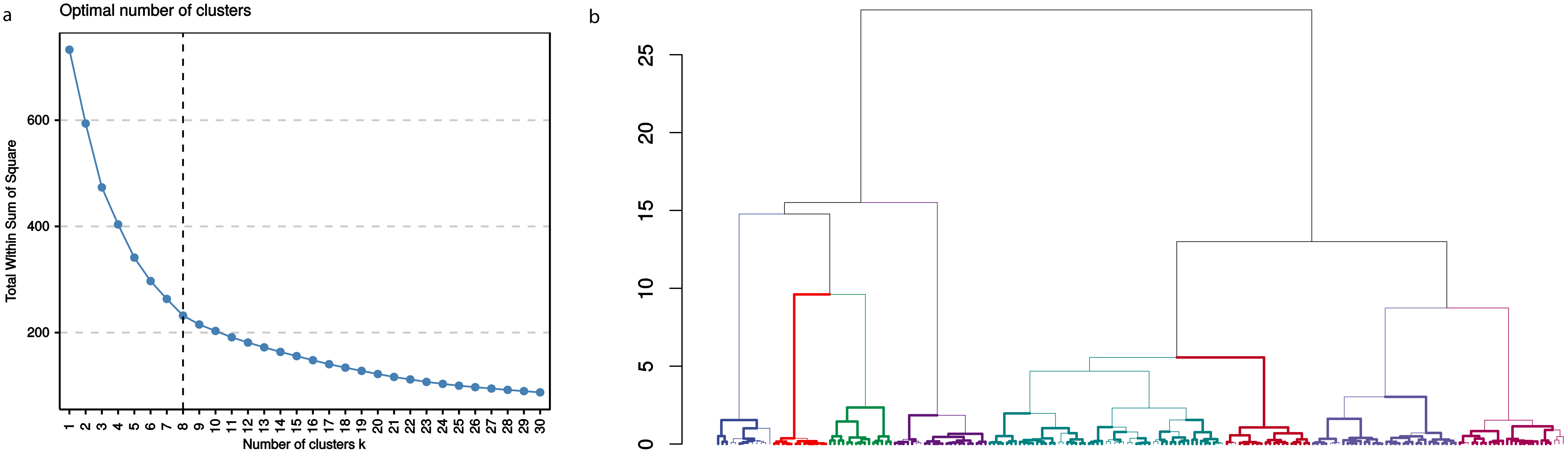

**Supplementary figure 10 - Clustering QC**

(a) Clustering estimation using optimum total within-cluster sum of square (wss). The final selection of the most optimal number of clusters was based on the knee in the blue line, e.g. the moment when increasing the number of clusters does not dramatically decrease wss.

(b) Bootstrapping results (5000 iterations) of the unsupervised hierarchical clustering with additional coloring of the eight defined clusters as used in this manuscript. Branches with Approximately Unbiased p-values (AU) ≤ 0.05 are highlighted with bolder lines and reflect significantly robust groups of samples based on similar characteristics derived from WGS. Y-axis displays clustering distance (Pearson correlation; ward.D).

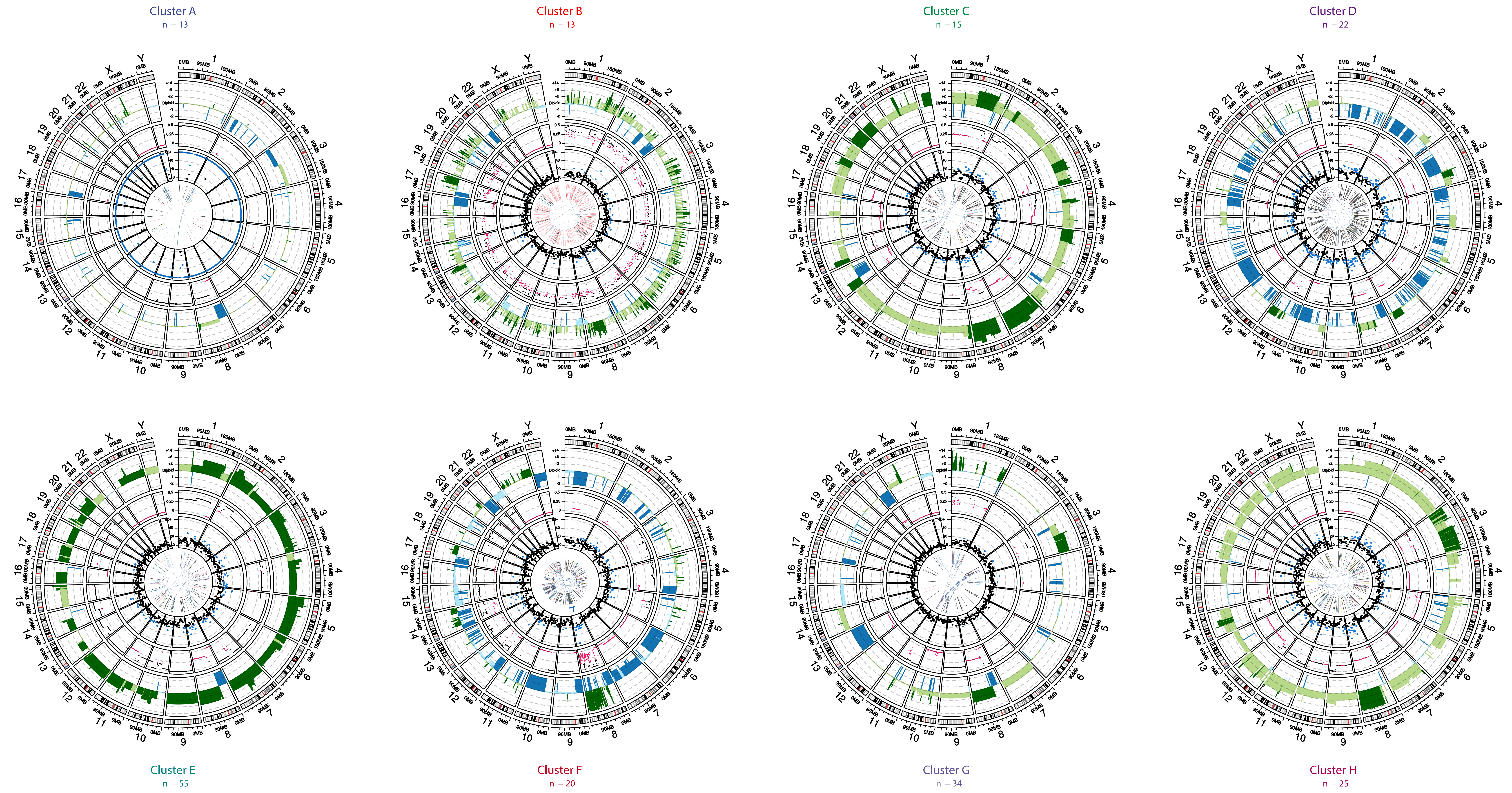

**Supplementary figure 11 - Representative mCRPC sample per cluster**

Genomic overviews of a single representative sample per (unsupervised) cluster. The outer track displays the genomic ideogram, the second-outer track displays copy number profiles (amplification in light green; deep amplification beyond sample-specific threshold (GISTIC2) in dark green, deletions in blue; deep deletions beyond sample-specific threshold (GISTIC2) in dark blue). The third track displays TC%-corrected minor allele-frequency (MAF) values of individual copy number segments (MAF ≤ 0.33 in pink; MAF ≥ 0.33 in black). The fourth track displays the number of mutations per 5 Mbp, ranging from 0 to 60+; bins with ≥ 20 mutations are highlighted in blue. The innermost track displays structural variants; interchromosomal translocations in dark blue, deletions in grey, insertions in yellow, inversion in light blue and tandem duplications in red.

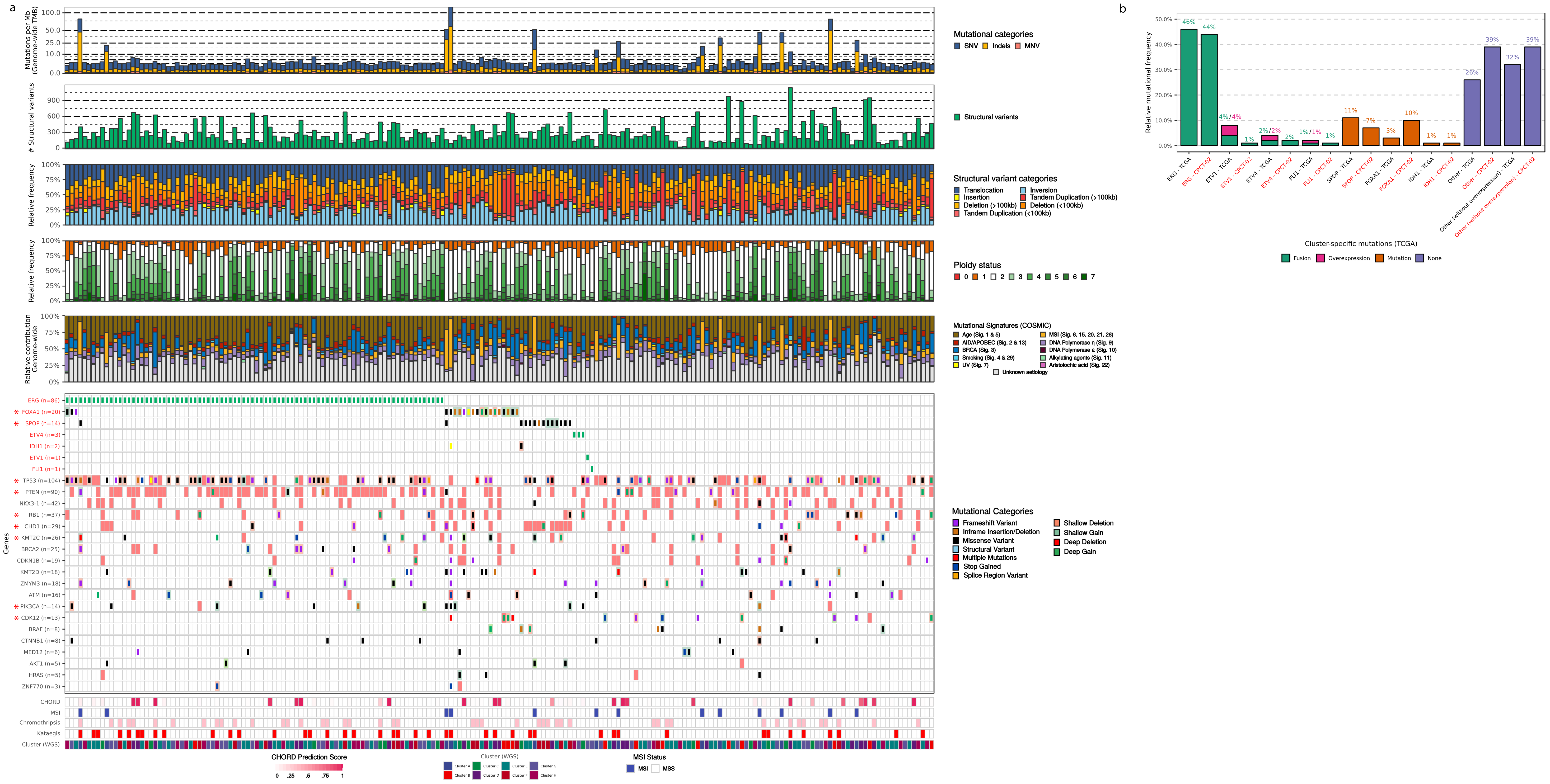

**Supplementary figure 12 - Supervised clustering of mCRPC based on TCGA criteria**

(a) Samples are sorted based on mutual-exclusivity of the same genes and aberrations as the TCGA clustering, depicted in red colors in the heatmap. In addition, all genes which the TCGA defined as recurrent alterations in primary prostate cancer are shown, genes also discovered in the mCRPC cohort as enriched in non-synonymous mutations or copy-number alterations (dN/dS and/or GISTIC2) are depicted with a red asterisk. The upper track displays the number of genomic mutations per Mbp (TMB) of SNV (blue), InDels (yellow) and MNV (orange) categories. Second track displays the absolute number of unique structural variants per sample. Third track displays the relative frequency per structural variant category, Tandem Duplications and Deletions are subdivided into > 100 kbp and <100 kbp categories. The fourth track displays the relative genome-wide ploidy status, ranging from 0 to ≥ 7 copies and the fifth track displays the relative contribution to mutational signatures (COSMIC) summarized per proposed etiology. The heatmap displays the type of mutation(s) per sample; (light-)green or (light-)red backgrounds depict copy number aberrations whilst the inner square depicts the type of (coding) mutation(s). In addition, the lower tracks display CHORD prediction score (HR-deficiency) (pink gradient), MSI status (blue), chromothripsis (pink), presence of kataegis (red) and in which of the eight genomic cluster, as defined by this manuscript, each sample falls.

(b) Overview of the relative frequency of samples captured per mutually-exclusive group for both the TCGA and mCRPC cohort. Promiscuous ETS family fusions (ETV1, ETV4 and FLI1) which were captured in the TCGA cohort using mRNA overexpression were split as the mCRPC cohort did not have accompanying mRNA sequencing data to perform a similar capturing.

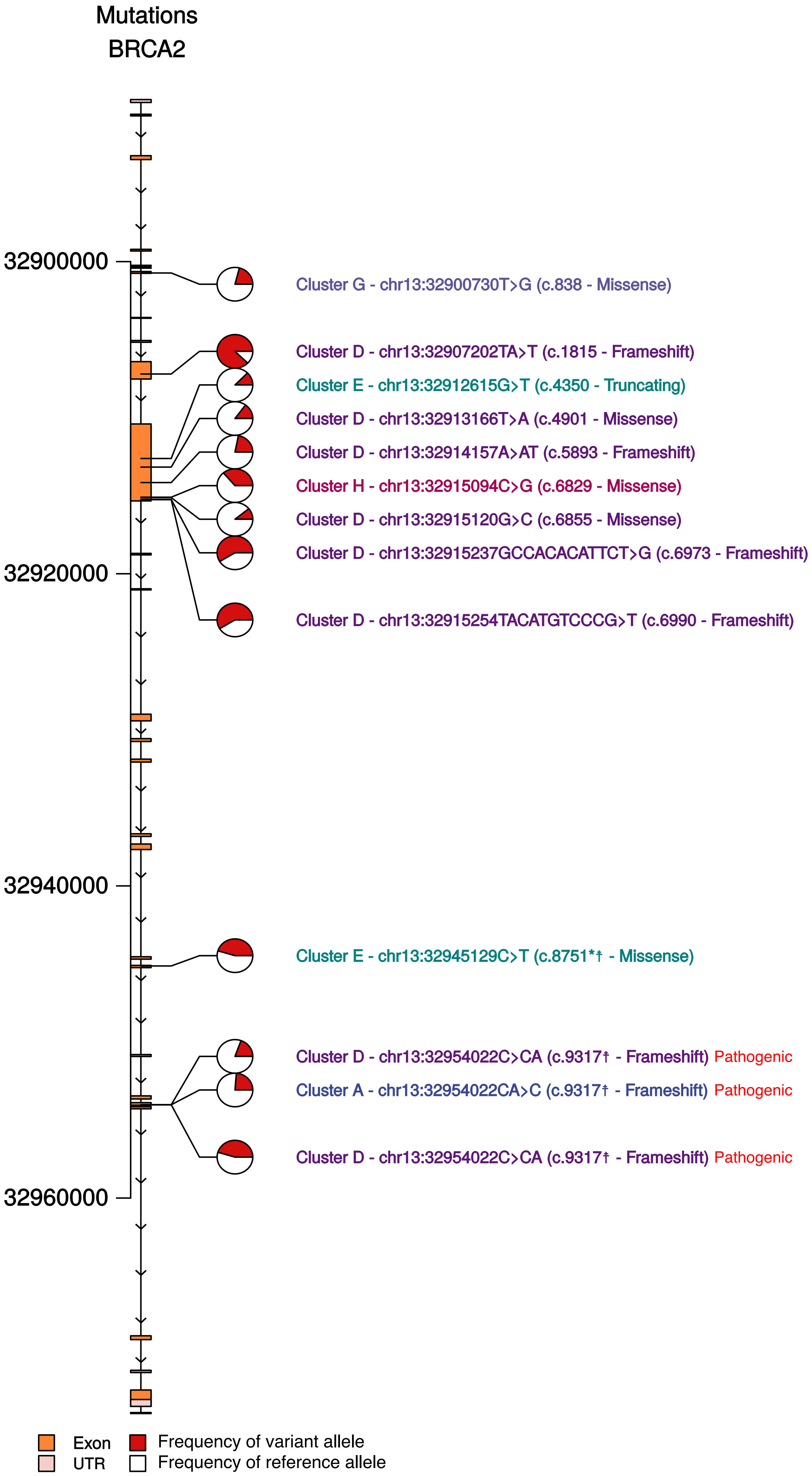

**Supplementary figure 13 - Overview of BRCA2 mutations**
Genomic distribution of non-synonymous BRCA2 mutations found within our mCRPC cohort. Mutations are displayed as pie-charts depicting the variant allele frequency (red portion of the pie-chart) and reference allele frequency (white portion of the pie-chart). Samples are colored based on their respective cluster after unsupervised clustering (figure 4).

Known COSMIC mutations are annotated with * and/or known dbSNP variants with a †. Alleles with known pathogenicity within ClinVar are highlighted, mutations without ClinVar annotation could be considered as variants with as-of-yet uncertain significance.

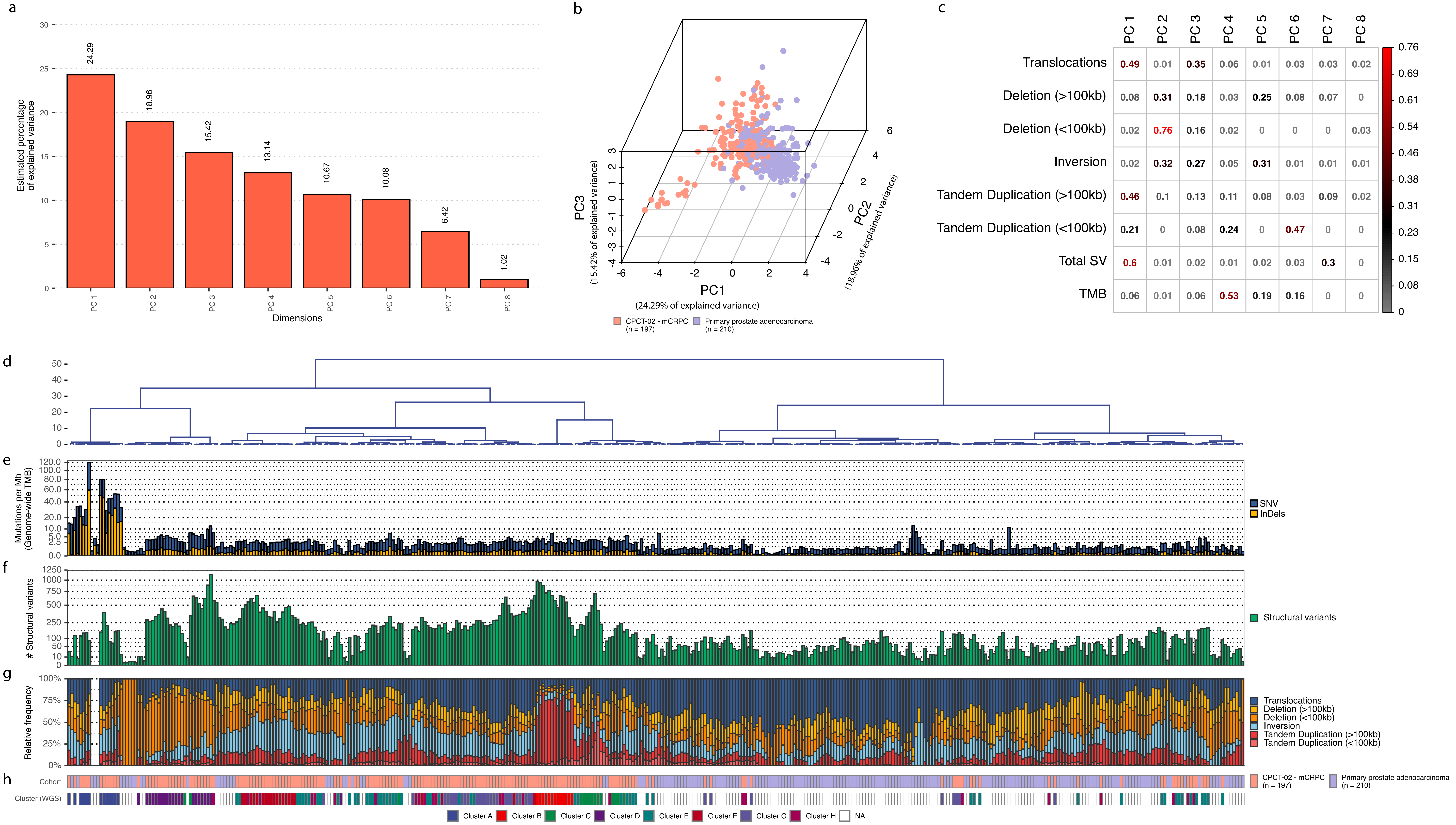

**Supplementary figure 14 - Overview of clustering scheme on primary prostate cancer and mCRPC**

(a) Overview of the explained variance per principal component (PC) in PCA of the combined dataset of primary prostate cancer (n = 210) and mCRPC (n = 197). PCA was performed in the following features: Total number of SV, genome-wide TMB (SNV and InDels) and relative frequency of structural variants (except insertions).

(b) Visualization of the first three principal components of PCA, each sample is colored based on their respective disease-setting, primary prostate cancer is colored as violet whilst mCRPC is colored as light-red.

(c) The quality of representation for each feature per principal component (cos2), this ranges from 0 (no importance / representation in PC) to 1 (absolute importance / representation in PC). Color gradient (0 to 0.7) denotes cos2, red values denote important / representation of feature within PC. Numbers shows are the cos2 values.

(d) Dendrogram of unsupervised clustering with optimal leaf ordering on genomic features. Y-axis displays clustering distance (Pearson correlation; ward.D).

(e) All genome-wide somatic SNVs (blue) and InDels (yellow) per Mbp (square root scale).

(f) Absolute frequency of structural variants per sample (square root scale).

(g) Relative frequency per structural variant category, Tandem Duplications and Deletions are subdivided into > 100 kbp and < 100 kbp categories.

(h) Respective cohort of the samples, primary prostate cancer is colored as violet whilst mCRPC is colored as light-red.

(i) Assigned clusters of mCRPC samples based on unsupervised clustering of genomic features as described in figure 4.

| Supplementary table 1: Participating centers | | |
|---|---|---|
| **Organization** | **Local principal investigator** | **Included patients for this study (n)** |
| Radboud UMC, Nijmegen | Carla van Herpen | 91 |
| Erasmus MC, Rotterdam | Martijn Lolkema | 38 |
| Franciscus Gasthuis & Vlietland, Rotterdam | Paul Hamberg | 20 |
| NKI-AVL, Amsterdam | Neeltje Steeghs | 15 |
| Isala, Zwolle | Jan Willen de Groot | 3 |
| Martini Ziekenhuis, Groningen | Johan van Rooijen | 3 |
| Medisch Centrum Leeuwarden | Hiltje de Graaf | 3 |
| Maastricht UMC, Maastricht | Vivianne Tjan-Heijnen | 3 |
| Noordwest Ziekenhuisgroep, Alkmaar | Mathijs Hendriks | 3 |
| UMC Utrecht, Utrecht | Els Witteveen | 3 |
| Amphia Ziekenhuis, Breda | Bert Jan ten Tije | 2 |
| Reinier de Graaf Gasthuis, Delft | Annelie Vulink | 2 |
| Treant Zorggroep, Hoogeveen | Sophia van den Boogerd | 2 |
| Zuyderland Medisch Centrum, Geleen | Frans Erdkamp | 2 |
| ETZ Elisabeth, Tilburg | Laurens Beerepoot | 1 |
| Leids Universitair Medisch Centrum, Leiden | Hans Gelderblom | 1 |
| Maasstad Ziekenhuis, Rotterdam | Rineke Leys | 1 |
| Meander Medisch Centrum, Amersfoort | Haiko Bloemendal | 1 |
| St. Antonius Ziekenhuis, Utrecht | Maartje Los | 1 |
| VUmc, Amsterdam | Henk Verheul | 1 |
| ZGT, Almelo | Esther Siemerink | 1 |

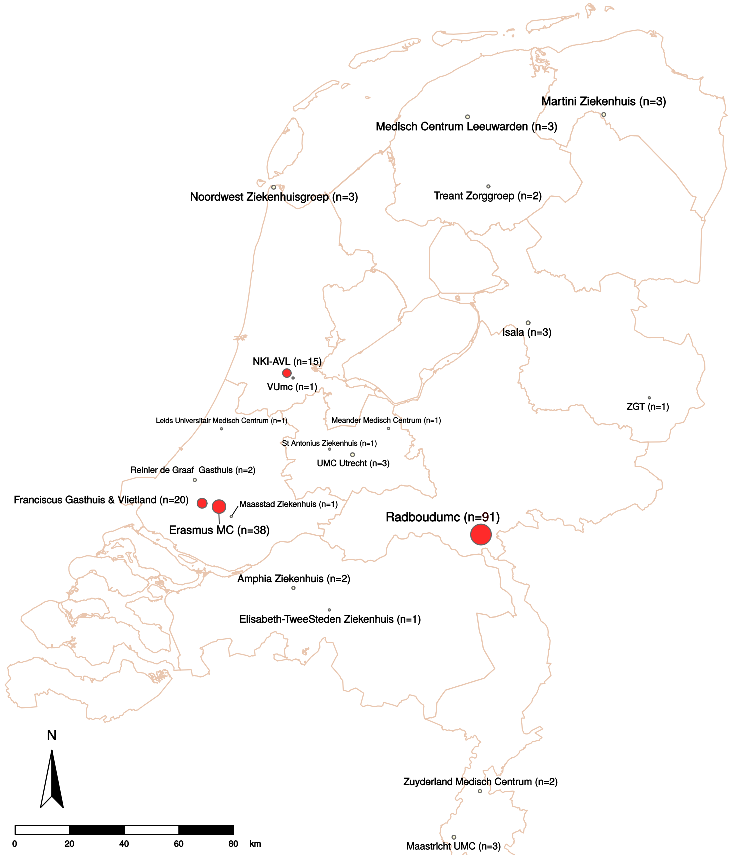

CPCT-02 (mCRPC)

| Supplementary table 2: Patient characteristics | | |
|---|---|---|
| *Patients (n=197)* | | |
| | n | % |
| **Age at biopsy** | | |
| Median | 68 | |
| Range (min-max) | 48-83 | |
| **Prior ADT** | | |
| Yes | 197 | 100,0 |
| Drug-based | 181 | 91,9 |
| Surgery-based (orchiectomy) | 3 | 1,5 |
| With Docetaxel | 6 | 3,0 |
| No clear documentation of ADT type | 7 | 3,6 |
| **Prior systemic therapy (other than ADT)** | | |
| 0 previous treatments | 27 | 13,7 |
| ≥ 1 previous treatments | 170 | 86,3 |
| 1 previous treatment | 45 | 22,8 |
| 2 previous treatments | 69 | 35,0 |
| 3 previous treatments | 31 | 15,7 |
| 4 previous treatments | 19 | 9,6 |
| 5 previous treatments | 6 | 3,0 |
| **Type of prior systemic therapy (other than ADT)** | | |
| Hormonal therapy only | 20 | 10,2 |
| Chemotherapy only | 37 | 18,8 |
| Radionucleotide therapy only | 4 | 2,0 |
| Immunotherapy only (Dendritic cell therapy) | 4 | 2,0 |
| Targeted therapy only | 0 | 0,0 |
| Hormonal and chemotherapy | 68 | 34,5 |
| Hormonal and radionucleotide therapy | 3 | 1,5 |
| Chemotherapy and radionucleotide therapy | 3 | 1,5 |
| Hormonal and immunotherapy | 3 | 1,5 |
| Chemotherapy and immunotherapy | 3 | 1,5 |
| Hormonal, chemotherapy and radionucleotide therapy | 15 | 7,6 |
| Hormonal, chemotherapy and immunotherapy | 4 | 2,0 |
| Hormonal, radionucleotide and immunotherapy | 2 | 1,0 |
| Hormonal, chemotherapy and targeted therapy (Olaparib) | 2 | 1,0 |
| Hormonal, chemotherapy, radionucleotide and immunotherapy | 1 | 0,5 |
| Unknown at time of analysis | 1 | 0,5 |
| **Prior radiotherapy** | | |
| Yes (curative radiotherapy of the prostate and/or palliative radiotherapy of metastases) | 117 | 59,4 |
| No | 77 | 39,1 |
| Unknown at time of analysis | 3 | 1,5 |
| **Started therapy after biopsy for whole-genome sequencing** | | |
| Yes | 138 | 70,1 |
| Hormonal therapy | 53 | 26,9 |
| Chemotherapy | 56 | 28,4 |
| Radionucleotide therapy | 12 | 6,1 |
| Immunotherapy (Pembrolizumab) | 6 | 3,0 |
| Targeted therapy | 3 | 1,5 |
| Combinational therapy | 6 | 3,0 |
| Other* | 2 | 1,0 |
| No | 19 | 9,6 |
| Unknown at time of analysis | 40 | 20,3 |
| **Biopsy site** | | |
| Liver | 29 | 14,7 |
| Lymph node | 81 | 41,1 |
| Bone | 70 | 35,5 |
| Lung | 3 | 1,5 |
| Soft tissue/Other** | 14 | 7,1 |
| *Boneregulating agent | | |
| **Soft tissue/other: (sub)cutis, muscle, peritoneum, kidney, bladder, adrenal gland | | |

**Supplementary data file 1 - Data used in figures**

Sheet A - Clinical characteristics per mCRPC patient.

Sheet B - Genomic characteristics per mCRPC patient.

Sheet C -Overview of detected kataegis foci.

Sheet D - Overview of kataegis foci characteristics.

Sheet E - Overview of predicted fusions genes.

Sheet F - Overview of predicted chromothripsis events.

Sheet G - Sample categorization into unsupervised clusters (A-H).

Sheet H - Mutually exclusive somatically mutated genes per clusters.

Sheet I - Peaks detected by GISTIC2 and the type of CN aberration per sample.

Sheet J - *AR*-enhancer and *AR* locus copy numbers per sample.

Sheet K - dN/dS driver-gene discovery output based on somatic mutations.

Sheet L - Mutational frequencies of TCGA-based clustering events.

Sheet M - Comparison of mutational frequencies of top 200 mCRPC-mutated genes (dN/dS, GISTIC2, subtype-specific and supplemented with top mutated genes based on number of total aberrations)

Sheet N - Comparison of mutational frequencies of top 200 mCRPC-mutated genes (dN/dS, GISTIC2, subtype-specific and supplemented with top mutated genes based on number of coding mutations)
